# Intestinal stem cell aging at single-cell resolution: functional perturbations alter cell developmental trajectory reversed by gerotherapeutics

**DOI:** 10.1101/2022.08.31.505869

**Authors:** Jiahn Choi, Michele Houston, Ruixuan Wang, Kenny Ye, Wenge Li, Xusheng Zhang, Derek M. Huffman, Leonard H. Augenlicht

## Abstract

The intestinal epithelium is consists of cells derived from continuously cycling Lgr5^hi^ intestinal stem cells (Lgr5^hi^ ISCs) that mature developmentally in an order fashion as the cells progress along the crypt-luminal axis. Perturbed function of Lgr5^hi^ ISCs with aging is well documented but the consequent impact on overall mucosal homeostasis has not been defined. Using single-cell RNA sequencing, the progressive maturation of progeny was dissected in mouse intestine, which revealed that transcriptional reprogramming with aging in Lgr5^hi^ ISCs retarded cell maturation in their progression along the crypt-luminal axis. Importantly, treatment with metformin or rapamycin at a late stage of mouse lifespan reversed the effects of aging on function of Lgr5^hi^ ISCs and subsequent maturation of progenitors. The effects of metformin and rapamycin overlapped in reversing changes of transcriptional profiles, but were also complementary, with metformin more efficient than rapamycin in correcting the developmental trajectory. Therefore, our data identify novel effects of aging on stem cells and the maturation of their daughter cells contributing to the decline of functional Lgr5^hi^ ISCs and the correction by geroprotectors.

## Introduction

The intestinal mucosa is a prime example of the decline in regenerative capacity, increase in cellular damage, and metabolic dysregulation that characterize aging (Di Giosia et al., 2022). These changes are linked to age-associated intestinal disorders, with aging also a major risk factor for colorectal cancer (CRC) (Nalapareddy, Zheng, & Geiger, 2022). The potential role of perturbations in Lgr5^hi^ intestinal stem cells (Lgr5^hi^ ISCs) are of particular interest in understanding aging since these are the cells that renew and maintain the mucosa through their continuous division at the crypt base (Barker et al., 2007). Reduction of Wnt signaling has been reported as a key that represses stem cell functions of Lgr5^hi^ cells with aging (Nalapareddy et al., 2017; Pentinmikko et al., 2019), but how it links to the developmental progression of progenitor compartments has not been addressed.

Several pharmacologic approaches have been reported to delay age-related manifestations including reversal of stem cell dysfunction and extension of lifespan (Moskalev, 2020; Partridge, Fuentealba, & Kennedy, 2020). Although none are yet approved for ameliorating effects of aging in humans, metformin and rapamycin have been studied (An et al., 2020; Blagosklonny, 2019; Moskalev et al., 2022; Novelle, Ali, Dieguez, Bernier, & de Cabo, 2016) as regulators of cellular metabolic activities that are linked to aging phenotypes. However, given the pleiotropic impact on metabolism and linked pathways, understanding the complexity of their effects requires more detailed investigation.

Therefore, we interrogated the complexity of aging effects on the intestinal mucosa and how potent geroprotectors intervene in the effects of aging using single cell RNA sequencing (scRNAseq) of the intestinal epithelium. Novel cellular and bioinformatics analyses revealed that aging significantly reprogrammed transcriptional profiles of canonical Lgr5^hi^ ISCs and delayed progression of developmental trajectory of the cells that were produced in the stem cell compartment. Further, heterogeneity of cellular response was identified that determines the efficiency of potential reversal effects by the geroprotectors, metformin and rapamycin. The data are incisive in defining how intestinal stem cells and the developmental programs of their progeny are altered with biological aging, identifying key imbalances among the pathways of Wnt signaling, cell cycling, and metabolism as cells undergo stage specific developmental maturation, and the overlapping but also distinct effects of metformin and rapamycin in correcting these defects in aged mice.

## Results

### Aging impairs function of canonical Lgr5^hi^ ISCs

Cell programming and function across all cell types and lineages of the intestinal mucosa were dissected in four groups of CB6F1 hybrid male mice: 5-month-old young mice (Y), 24-month-old aged mice (O) fed a purified diet for 3 months beginning at 2 months or 21 months of age, respectively, with two additional groups of aged mice also receiving the diet supplemented with either 0.1% metformin (O-met) or rapamycin (O-rap) at 42ppm for the 3 month period (Figure 1a). FACS isolated intestinal epithelial cells (Epcam+, Cd45neg) were analyzed by scRNAseq. Known markers aligned 21 cell clusters with cell types and lineages of the small intestine from stem through progenitor cell compartments and then less mature and eventually fully differentiated absorptive and secretory cells (Figure 1b, Figure S1a). The proportion of cells in each cluster were similar across the different age/treatment groups, suggesting that the overall cellular composition of the intestinal epithelium was maintained for > 2 years of age in the mouse (Figure S1b). However, analysis of gene expression demonstrated a wide range of genes altered in expression with age across clusters and lineages (≥ 1.5-fold change, adj *P* < 0.01, Figure 1c), suggesting that transcriptomic profiles in the ostensibly same cell types are reprogrammed with age. This was most prominent for fully mature enterocytes (EC8), which also increased in relative abundance in old mice (Figure 1c, Figure S1b). For most clusters, treatment of the aged mice with metformin or rapamycin reversed altered expression profiles in cells. Restoration of expression profile was most effective in EC8 cells, mature enterocytes, for both drugs, where it had been altered to the greatest extent in the aged mice (Figure 1c, Table S1).

**Figure 1:**
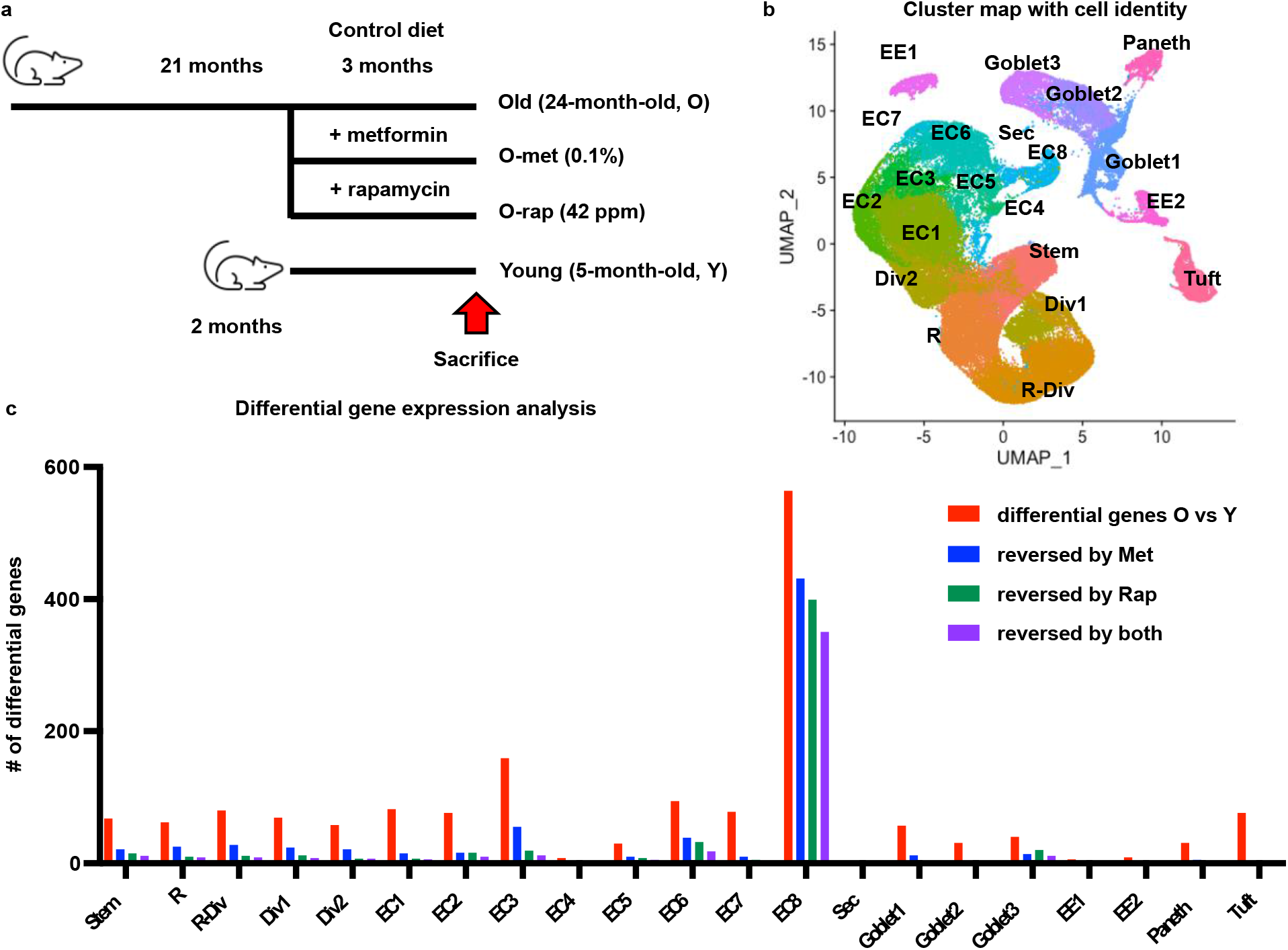
scRNAseq analysis of small intestinal epithelial cells: **a**, Schematic representation of experimental designs; drug treatment was started at age of 21-month-old for 3 months. **b**, Cluster map and cell lineages from 12 combined scRNAseq libraries. **c**, Number of differentially expressed genes (fold change > 1.5 & adj *P* < 0.01) in each cluster from comparison between old vs young (red bar), and number of genes reversed by metformin (blue bar), rapamycin (green bar) or both (violet bar).

The Lgr5 gene, encoding the receptor for the key stem cell growth factor R-spondin, is regulated by Wnt signaling, and in the intestine, is most highly expressed in cells at the crypt base (Barker et al., 2007). These Lgr5^hi^ cells are considered the major canonical stem cells maintaining structure and function of the intestinal epithelium (Barker et al., 2007). The mean expression of Lgr5 mRNA in the stem cluster from old mice significantly decreased by 33% compared to that in young mice, and was restored by metformin or rapamycin (Figure 2a). We next focused on Lgr5^hi^ cell signature genes, defined as those genes altered in expression in the immediate Lgr5^lower^ progeny of Lgr5^hi^ cells that have left the stem cell niche and no longer function as stem cells (Munoz et al., 2012). Of the 467 transcripts in this stem cell signature gene set expressed in Lgr5^hi^ cells from young mice, 71% were expressed at a lower level in older mice (Figure 2b).

**Figure 2:**
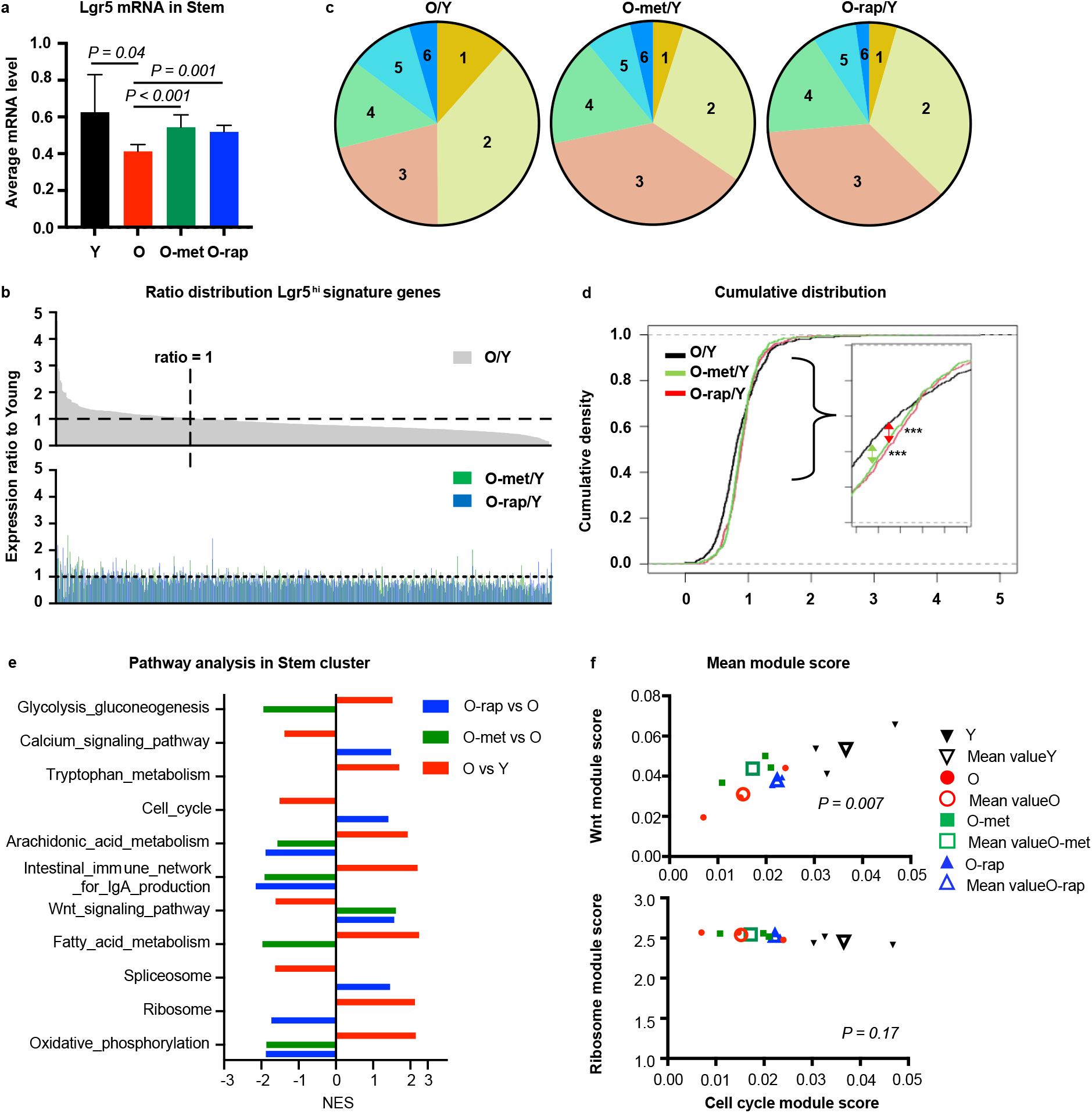
Impact of aging on function of intestinal stem cells: **a**, average mRNA expression of the *Lgr5* genes in the Stem cluster. **b**, Expression ratio of old vs young of 467 genes of an Lgr5^hi^ ISC signature (Munoz et al., 2012), and corresponding ratio of old-met vs young or old-rap vs young (lower panel). **c**, Distribution of genes as a function of ratio of expression; corresponding ratios in Table 1. **d**, Cumulative density graph of ratio distribution. Statistical analysis was performed using Kolmogorov–Smirnov test between two groups (***: *P* < 0.001). **e**, Differentially-regulated pathways (adj *P* < 0.05) in the Stem cluster using GSEA KEGG pathways; NES: normalized enrichment score. **f**, Scatter plot for two variables with mean module score. Mean module score from each mouse (filled) or each condition (empty, average value) were plotted for Wnt and cell cycle pathways (top) or Ribosome and cell cycle pathways (bottom). *P* value was calculated using MANOVA, assuming each mouse as independent.

The distributions of expression levels in young vs old mice, and in young vs old treated with either rapamycin or metformin, are shown in Figure 2c, with the percentage of genes at different levels of expression relative to young mice in Table 1. Compartments 1 and 2 are those Lgr5^hi^ stem cell signature genes repressed by >50% (compartment 1) or 20-50% (compartment 2, Figure 2c). However, for older mice treated with rapamycin or metformin, this pattern was shifted (Figure 2c, Table 1). Specifically, in old mice, 12% of the signature genes were expressed at or less than 50% of their level in young mice (compartment 1), but this decreased to 5% and 4% for the old mice treated with metformin and rapamycin, respectively. Similarly, expression level for 38% of the signature genes were 20% to 50% lower in old mice (compartment 2), but the percent of signature genes at these lower levels decreased to 30% and 33% for the old rapamycin and metformin treated mice, respectively. Reflecting this, the number of genes in compartments 3 and 4, in which expression level changes were within 20% of young mice, increased in the old mice treated with rapamycin or metformin (Figure 2c). Cumulative distribution of gene expression ratio was further analyzed using the Kolmogorov–Smirnov test (Massey Jr., 1951). Comparison between Old/Young vs Old-treatment/Young for each drug significantly shifted the distribution closer to 1 (*P* < 0.001 for each drug). However, comparison between rapamycin and metformin was not significantly different (Figure 2d). Thus, the repressed expression of stem cell signature genes in aged Lgr5^hi^ cells was significantly rescued by treatment with rapamycin or metformin.

**Table 1.** **% Lgr5 signature genes in range**

|  | **Expression ratio** | **O/Y** | **O-met/Y** | **O-rap/Y** |
| --- | --- | --- | --- | --- |
| **1** | **0 – 0.5** | **12%** | **5%** | **4%** |
| **2** | **0.5 – 0.8** | **38%** | **30%** | **33%** |
| **3** | **0.8 – 1** | **21%** | **37%** | **36%** |
| **4** | **1 – 1.2** | **14%** | **17%** | **17%** |
| **5** | **1.2 – 1.5** | **10%** | **7%** | **7%** |
| **6** | **> 1.5** | **4%** | **4%** | **2%** |

Gene set enrichment analysis (GSEA) identified significantly altered pathways in the stem cell cluster (Figure 2e). Oxidative phosphorylation (OXPHOS), a key metabolic pathway in regulating stem cell function (Rodriguez-Colman et al., 2017; Zhang et al., 2022) was upregulated in the stem cell cluster of older mice, and this was reversed by either metformin or rapamycin. Moreover, the linked metabolic pathways of fatty acid metabolism and glycolysis were also upregulated in the stem cluster of old mice, but reversed only by metformin treatment. In contrast, Wnt and cell cycle pathways, which are fundamental for stem cell self-renewal and generation of progeny that undergo differentiation, were suppressed in the stem cluster of old mice, while the ribosome pathway, also important for supporting stem cell function, and down-regulated as cells mature during their migration along the crypt-villus axis (Mariadason et al., 2005) was upregulated in the stem cells. Thus, although it is established that OXPHOS is necessary for ISC functions (Rodriguez-Colman et al., 2017), the elevation of this and related fatty acid metabolism in older mice may also perturb stem cell functions, emphasizing the importance of maintaining these pathways within a homeostatic range for normal stem cell functions. This was confirmed by investigating the self-renewal capability of stem cells by analysis of BrdU incorporation prior to sacrifice (Figure S2a). Quantification of BrdU+ nuclei in duodenal crypts confirmed that proliferating cells were reduced in old mice by 28% (*P* <0.0001; Figure S2b). Metformin or rapamycin treatment increased proliferating cells back to youthful levels, and both reached statistical significance (Figure S2b), confirming that proliferation of ISCs was suppressed both transcriptionally and functionally, which were rescued by drug treatments.

Single-cell analysis provides the ability to investigate the link between the essential and altered pathways in individual cells. This was investigated using the AddModuleScore function which calculates activity score for a pathway in individual cells by accessing gene expression level compared to random control genes (Tirosh et al., 2016). Supplementary Figure 2c illustrates correlated distribution of Wnt and cell cycle pathways in individual ISCs (left) and Lgr5 ^hi^ ISCs (right) for each condition, the elliptical areas delineating points within 95% of the Gaussian distribution. This shifted towards the lower left with aging, but was partially reversed by rapamycin and metformin back towards the informatic space occupied by younger mice. In contrast, coordinate regulation of the ribosome (protein synthesis) and cell cycle pathways showed that the module score was higher for ribosome activity with lower cell cycle activity in aged mice, but the effect of either rapamycin or metformin treatment to shift these back towards the pattern of young mice were less clear (Figure S2d). Bivariate analysis using MANOVA for two variables based on mean module score showed that the Wnt and cell cycle pathways were positively correlated and significantly different among all experimental groups, demonstrating that Wnt and cell cycle pathways are coordinated in individual cells and average activities in the stem cluster shifted significantly among groups (*P* = 0.007). However, ribosome and cell cycle pathways were not clearly coordinated among all groups (Figure 2f). Thus, for cells in the ISC compartment overall, and specifically for the Lgr5^hi^ ISCs, the Wnt and cell cycle pathways are both regulated coordinately, significantly suppressed by aging, but enhanced ribosomal activity in aged mice is discordant with the repressed cell cycle pathway, suggesting perturbed balance between protein synthesis and cell cycle in aged ISCs.

### Altered metabolism of ISCs influence their essential functions

The perturbed balance between self-renewal and differentiation for the Lgr5^hi^ cells, and the fact that metabolic changes of ISCs influence how their progeny mature and undergo lineage differentiation (Alonso & Yilmaz, 2018; Beumer & Clevers, 2020; Chandel, Jasper, Ho, & Passegue, 2016), extended analyses to how these cells mature as they progress through progenitor and lineage-specific compartments. Pathway analysis using fold change of differentially expressed genes in old vs young mice showed that at least 79% of the significantly altered pathways in ISCs of old mice continue to be altered in the progenitor cell types sequentially emerging from the stem cell compartment (Figure S3a). This included the compartments identified as Replicating (R), Replicating-Dividing (R-Div) and Dividing 1 and 2 (Div1, Div2). Therefore, we hypothesized that changes in stem cell expression profile can continue to influence the progressive maturation of cells as they progress through these cell compartments. This was analyzed using Monocle, which predicts trajectory of cell maturation based on gene expression profiles in individual cells (Trapnell et al., 2014). Assuming cells derived from the Stem cluster (yellow arrow, Figure S3b), monocle established a trajectory from Stem through intermediate cell types towards enterocytes for the young mice (Figure S3b). Spatial markers that have been shown to identify cell position along the crypt-villus axis (Moor et al., 2018) confirmed that the predicted trajectory was well aligned with physical positioning of lineage development along the crypt-villus axis (Figure S3c), including the Monocle prediction of the dividing cells down-stream of the replicating cells. Similar trajectories were predicted for both young and old mice, irrespective of drug treatment. This is consistent with the data that cell-type representation was well conserved among all the groups (Figure S1b). To follow cell progression, cells were selected along the trajectory from the stem cell origin (yellow arrow) to the point at which the daughter cells have traversed the progenitor cell compartments to give rise to enterocytes (green arrow, Figure S3b, color-coded either green or pink in Figure 3a). Along this portion of the trajectory, there was a main developmental route and positions branching off along the route. To interrogate how cellular characteristics differed depending on their position along the trajectory, cells were dichotomized into mainstream (main, green color) and side branch cells (side, pink color) (Figure 3a). Module scores for the pathways identified in Figure 2e were calculated as previously described to compare the main and side branch cells of each mouse group (Figure 3b-i). Strikingly, violin plots of module scores clearly showed that the alterations induced by aging were principally derived from mainstream cells along the path of developmental cell maturation. For instance, distribution and average module score for Wnt, cell cycle, Ribosome and OXPHOS pathways from main cells were significantly different among groups (Figure 3b-e) but module scores of the same pathways for side cells did not differ (Figure 3f-i). ANOVA analysis identified that the mean module score for the Wnt and ribosome pathways were significantly changed with conditions for main stem cells, but not for side stem cells, suggesting that the impact of aging is intensified in cells that drive differentiation. Follow-up using Tukey’s multiple comparison test (Table S2) show the groups for which the difference was significant. Based on the adjusted P value in Table S2, the impact of aging for all three pathways was significant only for the main trajectory cells. Metformin treatment boosted Wnt and cell cycle activities in these main stem cells, which restored the pattern found in young mice. Rapamycin also elevated these pathways in the main stem cells of old mice, but only the cell cycle pathway was significantly reversed (Figure 3b-d, Table S2). This dichotomized analysis also revealed that OXPHOS activity differed based on cell position on the trajectory. In the main stem cells, there was a steep difference of mean module score between young and old mice, but the difference did not exist in the side stem cells, implying that aging impaired the balance of metabolic activity determining function of ISCs (Figure 3e, i). Elevation of OXPHOS in main stem cells from old mice was reversed towards that of young mice with metformin, but rapamycin was not effective for main stem cells, rather affecting side stem cells, reflecting the overall lower efficacy of rapamycin and the distinct mechanisms of action of the two drugs (Figure 3e, i). Follow-up analysis using Tukey’s multiple comparison after ANOVA confirmed that elevation of OXPHOS in main old stem cells was reversed with metformin treatment, but not with rapamycin (Table S2). Thus, this bioinformatic analysis shows that activities of key cellular functions and metabolic pathways are distinct in relation to position along the intestinal cell developmental trajectory in young mice, compared to old mice. Metformin and rapamycin treatment both augmented key pathway activities. However, metformin showed greater effect on adjusting key pathways in main cells, while rapamycin effect was less dependent on the cellular position along the trajectory.

**Figure 3:**
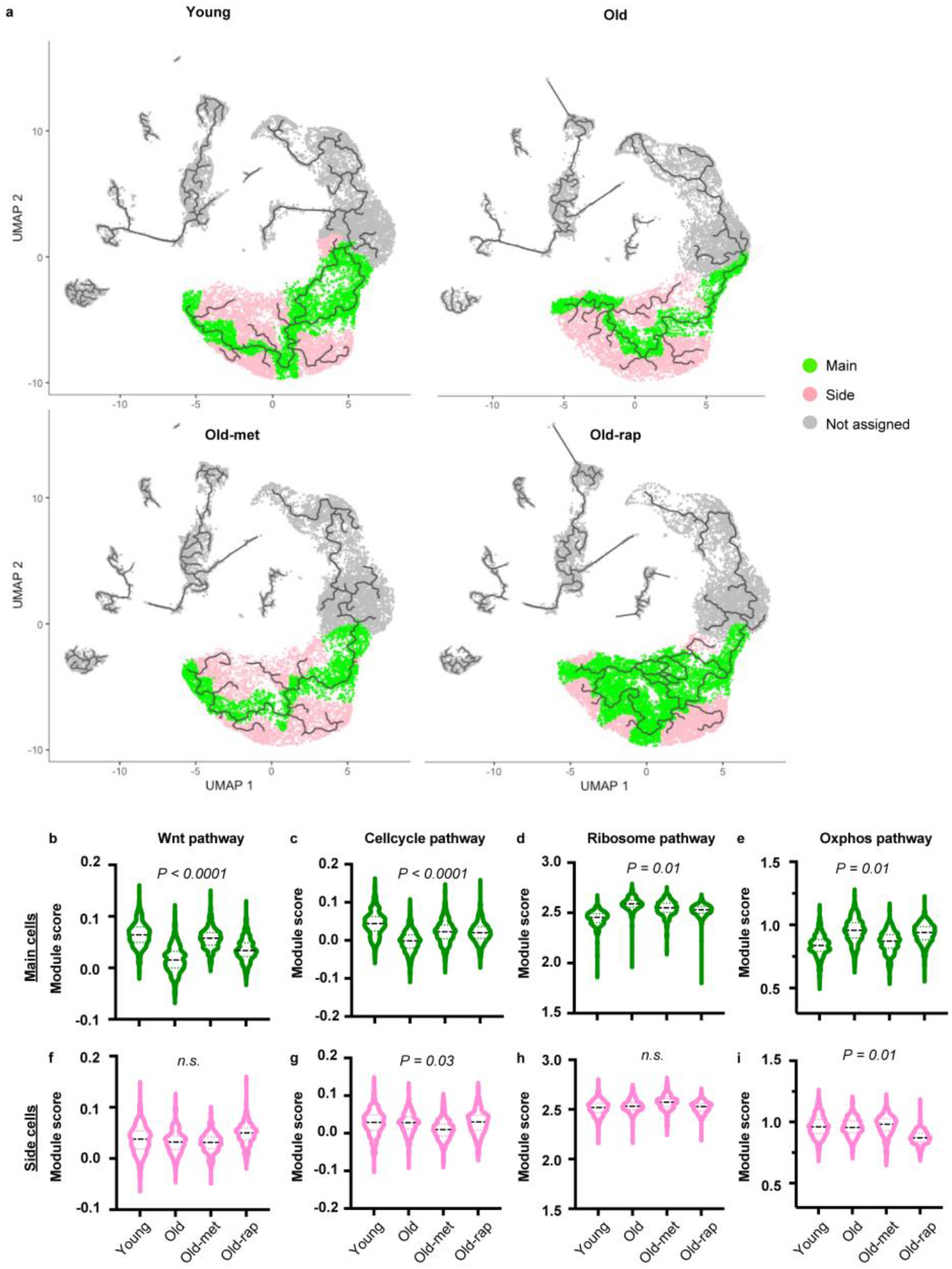
Impact of aging on stem cells differs depending on the position along the trajectory: **a**, trajectory graph for each condition. Based on position of each cell along the trajectory, cells on main track color-coded green, and cells on side branches color-coded pink. Cells not included in analysis are color-coded grey. **b-i**, module scores of four different pathways plotted separately by position (main: b-e, side: f-i) on the track and condition. Statistical significance was determined by ANOVA test and applied for each position separately.

### Alteration in stem cells delays proper maturation of cells

Analysis of disease pathways for the mainstream cells along the trajectory using Ingenuity Pathway Analysis (IPA) revealed that the stem and progenitor clusters along the trajectory in old mice showed activation of the predefined pathways of organismal death, growth failure, and apoptosis, and complementary inhibition of cell survival pathways, consistent with potential deteriorating function of the tissue with age (Figure 4a-d). These pathogenic pathways reversed in the stem and R clusters (early cell type along the developmental trajectory) of old mice by treatment with either metformin or rapamycin (Figure 4a-d). However, further downstream in the developmental trajectory (R-Div, Div-1, Div-2), efficacy of the two drugs diverged, with metformin significantly more effective in reversing the pathway changes with age (Figure 4a-d). Therefore, we extended the informatic analysis to how cell maturation was altered by aging and drug treatment throughout the developmental trajectory of the cells by interrogating cells on the branching points between sequential compartments in the trajectory. This was done for cells informatically identified as Stem, R, R-Div, Div1, Div2, and Others, the latter encompassing later differentiated cells (Figure 4e). Each of these cell types emerged sequentially in Young mice along the trajectory (Row 1, Figure 4e). However, in Old mice (Row 2), there was a clear delay along the trajectory in the development of each cell type. Strikingly, metformin treatment (Row 3) restored the temporal appearance of cell type to a more youthful position all along the trajectory. Rapamycin was similarly effective, with the exception of Div1 cells, which did not differ from Old mice (Row 4) in line with its lower efficiency on later cell types rescuing pathogenic pathways in Figure 4a-d.

**Figure 4:**
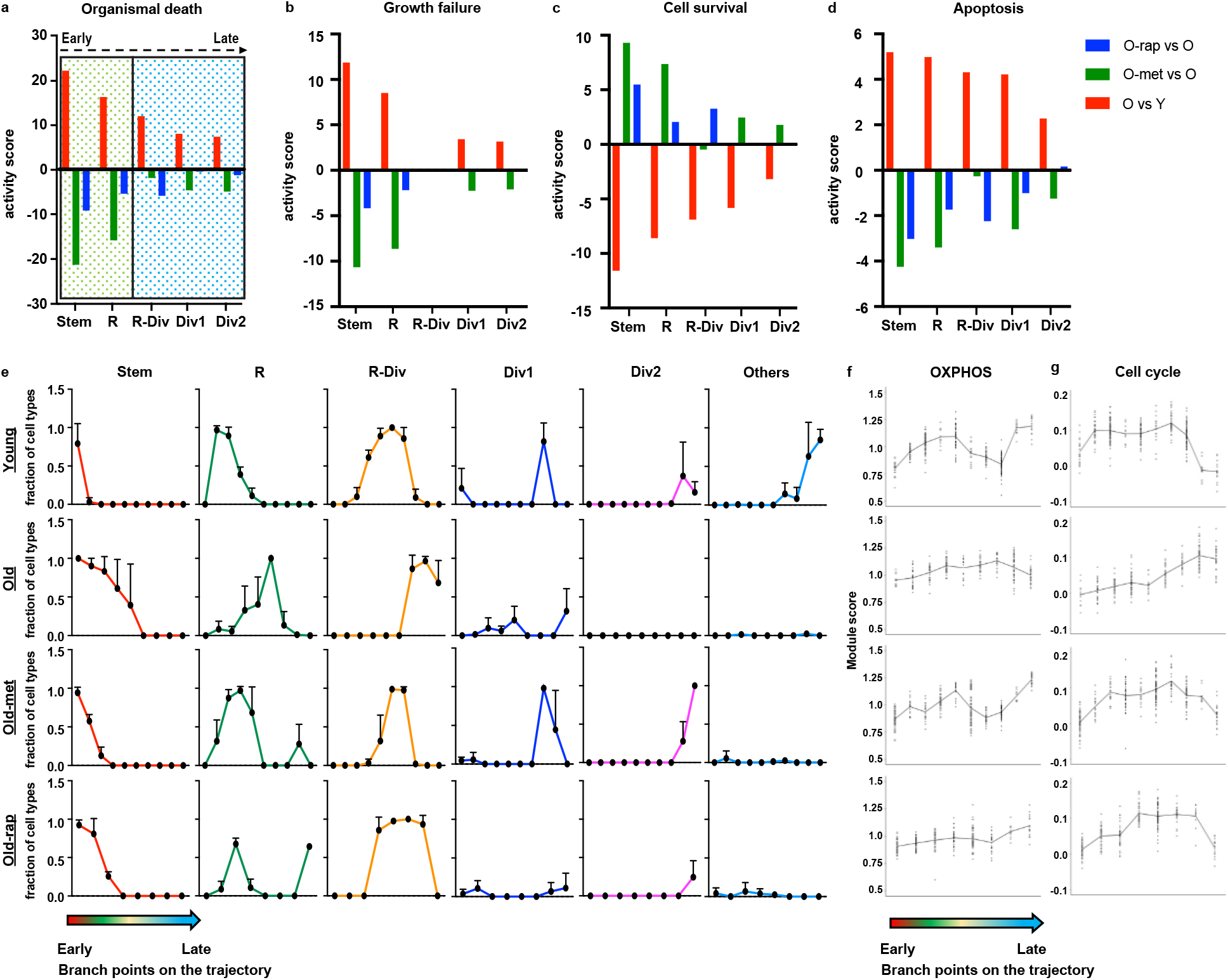
Aging delays proper cell maturation along the developmental trajectory: **a-d**, assignment of disease pathway based on differentially expressed genes (*P* adj < 0.05) in main cells of Stem and subsequent progeny clusters were analyzed using QIAGEN IPA (QIAGEN Inc., https://digitalinsights.qiagen.com/IPA). Dotted green box indicates cell types assigned to clusters early in the trajectory (Stem and R); dotted blue box indicates cell types in later in the trajectory (R-Div, Div1, and Div2). **e**, cell type transition along the branch points of the trajectory plot. **f**, Module score of OXPHOS pathway at each branch point analyzed for cell type transition in e. **g**, Module score of Cell cycle pathway at each branch point analyzed for cell type transition in e.

Finally, the relationship between the OXPHOS and cell cycle pathways was investigated along this trajectory using a module score for each pathway as in the earlier analyses. In young mice, the mean module score for OXPHOS between branch points (Figure 4f) gradually increased and then decreased along with the cell type transition of the developmental trajectory, and showed a sharper rise at the last 2 stages of the trajectory where mature cells initially appear. However, this pattern was greatly attenuated in Old mice. Metformin restored the pattern towards that of young mice, but rapamycin did not. Similarly, for the cell cycle pathway (Figure 4g), the mean module score increased as the cells emerged from the stem cell compartment, in parallel to known elevated proliferation in subsequent compartments, collectively termed the transit amplifying cells, and then fell sharply at the end of the trajectory in accordance with appearing of matured enterocytes. In contrast, there was a progressive increase in the cell cycle module score along the trajectory in Old mice, but the sharp decrease at the end of the trajectory seen for Young mice did not occur, suggesting an incomplete process of maturation until the last branch point on the trajectory. However, both metformin and rapamycin restored this overall pattern in old mice to more closely resemble the temporal pattern at these stages in young mice. In summary, our data suggest that cellular function is tightly linked to their metabolic activities driving proper maturation along the developmental trajectory, and aging perturbs precise coordination which leads to impaired cellular progression.

## Discussion

Transcriptomics at single cell resolution establish that aging suppressed expression of the Lgr5 gene in stem cluster by >30% and repressed expression of >70% genes reported as the transcriptional signature defining Lgr5^hi^ stem cells (Munoz et al., 2012). Lgr5 is a Wnt target gene and a receptor for R-spondin, a functional driver of cycling ISCs, and we documented down regulation of Lgr5 expression in line with repression of Wnt signaling pathways in stem cells of the aged mice. Therefore, aging may compromise sustained self-renewal and the ability of these canonical stem cells to produce progeny and maintain mucosal function. Elsewhere, we showed that the extensive down-regulation of the stem cell signature genes in Lgr5^hi^ cells begins as early as middle age (12-month-old mouse) in comparison to young adults (3-month-old) (Choi et al., 2022). Thus, these data support reports that aging impairs function of the primary stem cells in the small intestine both functionally and transcriptionally (Cui et al., 2019; Nalapareddy et al., 2017).

Metabolic pathways are critical for sustaining continuous cell cycling of Lgr5^hi^ ISCs to maintain the mucosa, and metabolic status is also a key in determining whether stem cells retain self-renewal capacity or differentiate following their replication (Mihaylova et al., 2018; Rodriguez-Colman et al., 2017; Wang, Odle, & Liu, 2021). Our data show that aged mice exhibited significant alterations in key metabolic pathways such as OXPHOS, glycolysis, and fatty acid metabolism. Therefore, metabolic programming is reconfigured in aging with important consequences for stem and progenitor cell functions and mucosal remodeling. Dichotomized analysis clearly showed that there was heterogeneity even among cells within the same cluster as regards to metabolic pathway profile. In young mice, stem cells exhibited lower expression of the OXPHOS machinery consistent with elevated cell cycle and Wnt pathway activities, while in old mice, OXPHOS was elevated in stem cells, in conjunction with suppression of Wnt and cell cycle pathways. Importantly, the unique architecture of the mucosa permits linking these alterations to developmental progression of the stem cell progeny. In this regard, the distinct metabolic activities linked to cell position along the crypt-luminal axis emphasizes that coordination between metabolism and stem cell functions is fundamental for normal stem cell functions, and that the disruption in aged mice alters the progression of cells along the mainstream of their developmental maturation. We have previously suggested this fundamental link is through altered mitochondrial function (Augenlicht & Heerdt, 2001), and more recently have shown this is a key to nutritional remodeling of ISCs and mucosal homeostasis that establishes elevated probability for development of sporadic tumors (Choi et al., 2022).

Metformin and rapamycin are gerotherapeutics under intense investigation for potential to delay many facets of aging to ameliorate physiological deterioration and potentially increase quality and length of life (Harrison et al., 2009; Martin-Montalvo et al., 2013; Novelle et al., 2016). Both pharmacological agents reversed multiple aspects of the aging transcriptomic phenotype in the intestinal mucosa. At doses shown to extend mouse lifespan, each drug suppressed OXPHOS and augmented Wnt signaling (Figure 2e). Interestingly, there were distinct differences in their impact on cell programming, with metformin more effective in adjusting metabolic pathway expression signatures by reducing the aging-associated effect on fatty acid oxidation and glycolysis while rapamycin more directly enhanced proliferation (Figure 2e). Importantly, further dissecting their distinct effects using dichotomized analysis revealed that the effect of metformin concentrated on main cells that drive developmental progression, while the effect of rapamycin is less linked to their position on the trajectory. Therefore, tracing cellular maturation using trajectory analysis showed that metformin-treated mice restored the profile of developmental progression that characterize the mucosa of young mice, while the effect of rapamycin was more limited. A recent study reported that intervention by inducing reprogramming factors in old mice showed rejuvenating effects that accompanied by restoration of metabolites at the systemic level (Browder et al., 2022), which supports our finding that correction of metabolic pathways was more effective on restoring phenotypic changes in old mice. Distinct mechanisms in the way two drugs reprogram cells suggests that combining the drugs may have complementary therapeutic effects in ameliorating the impact of aging.

Aging is a major underlying risk factor for CRC (Nalapareddy et al., 2022), with clear effects of aging on the intestine reported (Baron & Pisani, 2021; Choi et al., 2018; He et al., 2020), and herein deconvolved by the single cell analyses. Of note, despite a major disruption in stem cell function and development of linages, older wild-type mice do not develop tumors spontaneously when fed control diets. However, when wild-type mice are fed a diet long-term that mimics intake levels of common nutrients linked to those consumed in populations at high risk for colon cancer, they do develop small and large intestinal tumors reflecting the etiology, incidence, frequency and lag with older age of human sporadic colon cancer (Aslam, Paruchuri, Bhagavathula, & Varani, 2010; Newmark et al., 2009; Newmark et al., 2001; Yang et al., 2008). This emphasizes that long-term dietary patterns and aging are inextricably linked as major risk factors for development of human CRC. In the dietary model in which sporadic tumors do develop, there is suppressed function of Lgr5^hi^ ISCs with alternate *Bmi1+, Ascl2*^*hi*^ cells mobilized that remodel the mucosa and establish pro-tumorigenic chronic inflammation (Choi et al., 2022; Li et al., 2019; Newmark et al., 2009). While aging also had a major impact on suppressing Lgr5^hi^ ISCs, there was no evidence that Bmi1+ cells were mobilized or other possible alternative stem cells recruited. This raises the important question of how aging and long-term consumption of a highly relevant high-risk diet interact in altering mucosal stem cell functions and physiology in establishing high risk for colon cancer, a disease that dramatically rises in older individuals. Moreover, since geroprotectors such as metformin and rapamycin hold promise for mitigating age-related alterations to the intestinal epithelium, the potential interaction of such drugs with dietary interventions may need to be explored for efficacy in mitigating aging related intestinal stem cell programming and lineage differentiation, and in preventing disease that limits quality of life and lifespan.

## Methods

### Animals and experimental design

CB6F1 hybrid male mice were obtained at either 2 or 21 months of age from the NIA Aged rodent colony. On arrival, young (Y) mice were immediately switched to a purified control diet and old (O) mice were randomized to either the same control diet, or a diet in which either 0.1% metformin (Cayman Chem, Ann Arbor, MI) or microencapsulated rapamycin (eRapa) at a concentration of 42 ppm (Rapamycinholdings, Inc., San Antonio, TX) was incorporated into the formulation (TestDiet, St. Louis, MO). Eudragit was also included into the control and metformin diets (429ppm) in order to match the rapamycin formulation. All mice were group housed and fed these diets for 3 months under a 14L:10D photoperiod at 22°C, and remained on these formulations for 3 months, and were subsequently euthanized and intestinal tissue was harvested at either 5- or 24-months of age, respectively, for analysis. All experimental methods were approved by the IACUC at the Albert Einstein College of Medicine.

### Single cell RNAseq

Mice were sacrificed, excised intestines opened longitudinally, rinsed with cold saline, single cell suspensions prepared, re-suspended in antibody staining buffer, blocked with FcR Blocking Reagent (Miltenyi Biotec, 130-092-575), washed and pelleted, then incubated for 20 minutes at 4°C with, EpCAM-APC (Miltenyi Biotec, Cat No. 130-102-234) and CD45-PerCP (Miltenyi Biotec, Cat No. 130-102-469). Cells were sorted by FACs on a MoFlo instrument and Epcam positive, CD45 negative total epithelial cells collected.

scRNAseq libraries were constructed by the Albert Einstein Genomics Core using the 3’ kit protocols of 10X Genomics (San Francisco, CA) with approximately 10,000 single FACS collected epithelial cells from each mouse processed on a Chromium Chip B microfluidic apparatus (10X Genomics). Library quality was verified by size analysis of transcripts (BioAnalyzer; Agilent, Santa Clara, CA) and after passing quality control assays, multiple libraries were multiplexed and sequenced by HiSeq PE150 using pair-end sequencing, with a readout length of 150 bases (Novogene; Sacramento, California), and the data assigned to individual cells using the sample indexes.

For sequence alignment and subsequent analysis, output Illumina base call files were converted to FASTQ files (bcl2fastq), these from each mouse aligned to the mouse mm10 genome v1.2.0 and converted to count matrices (Cell Ranger software v3.0.2). 5000-8000 individual cells identified for each sample, and unique molecular identifiers (UMI) were used to remove PCR duplicates. Quality control and downstream analyses were done in R v4.1.2, using Seurat package v4.1.0(Stuart et al., 2019). To discard doublets or dead cells, cells with gene counts <200 or >5000, or a mitochondrial gene ratio>20%, were filtered out. Cells from different samples were integrated using Seurat FindIntegrationAnchors and IntegrateData functions, and clusters were identified from the integrated data set using the Seurat FindClusters function. This calculates k-nearest neighbors according to Principle Component Analysis, constructs a shared nearest neighbor graph, and then optimizes a modularity function (Louvain algorithm) to determine cell clusters. Cell clusters were identified using established cell-type markers for intestinal epithelial cells and cluster markers were identified using Seurat FindMarkers function. Differential gene expression compared samples among experimental groups: initial criteria were an expression change of ≥1.5 fold with associated adjusted P-value of <0.01 in group comparison (Seurat FindMarkers function). Pathway analysis was performed on differentially expressed genes using clusterProfiler R package v4.2.2; gene set enrichment analysis (GSEA) used the fgsea R package (v1.20.0), and the MSigDB (v5.2) KEGG pathway database. Pathways at Adj P <0.05 were considered statistically significant. Trajectory analysis was done using Monocle3 R package v1.2.6(Trapnell et al., 2014) with cluster information and cell-type identifications migrated from Seurat using an in-house developed code. The integrated dataset was divided among different experimental conditions, and trajectories were generated for each condition. Module score to determine average expression of gene set for each pathway was calculated with Seurat AddModuleScore function(Tirosh et al., 2016). Same gene list for KEGG pathway analysis was used.

### Immunohistochemistry for BrdU+ cells

Immunostaining was performed as previously described. Briefly, antigen retrieval was performed via a citrate buffer (pH=6) using a pressure cooker for 10 minutes. After rehydration and blocking endogenous peroxidase by 3.0% H2O2 for 5 min, avidin and biotin blocking (Vector Labs SP-2001) was performed each for 15 minutes with a wash step in between. Next, slides were blocked and stained using a mouse-on-mouse (M.O.M) kit (Vector Labs BMK-2202) with an antibody against BrdU (1:200; cat#5292, Cell Signaling) following the manufacturing’s protocol. Slides were then incubated with biotinylated secondary antibody for 30 min, followed by a streptavidin-HRP detection system (Vector labs PK-4000) and application of 3, 3′-diaminobenzidine (DAB) for visualization of the antigen-antibody complex (Scytek).

### Image analysis

To quantify BrdU+ nuclei, duodenal epithelium was imaged at 60x magnification. Automatically set threshold was applied to count BrdU+ nuclei. 15 crypts were measured from 3 individual mice in each group. Number of positive nuclei were divided by total nuclei in the crypt. One-way ANOVA followed by Tukey’s multiple comparisons test were applied to calculate *P* value. Statistical analysis was performed using GraphPad Prism version 0.3.1 for Mac, GraphPad Software, San Diego, California USA, www.graphpad.com.

### Statistical Analysis

Differences in gene expression between groups (Figure 2a) were modelled by a negative binomial mixed effect model with the number of read counts of individual cells as the response, offset by the number of the total counts of the cells, and the individual mice are treated as random effects. The computation is performed by R function glmer.nb. Two-sample Kolmogorov-Smirnov test was used to compare two distributions (in Figure 2d). Multivariate analysis of variance was used to compare mean module scores at two pathways among the four experimental groups (in Figure 2f) using R function MANOVA assuming each mouse as independent.

Figure 3b-i was tested using one-way ANOVA followed by Tukey’s multiple comparisons test (Table S2), which was performed using GraphPad Prism version 0.3.1 for Mac, GraphPad Software, San Diego, California USA, www.graphpad.com.

## Supporting information

included

## Acknowledgements

Supported in part by P30 AG038072, 5T32AG023475-20 from the NIA, and R01CA214625, R01CA229216, R01CA222358, and P30-013330 from the NCI.

## Author contributions

L.H.A., and D.M.H., designed experiments and supervised the work. K.Y performed statistical analysis. J.C. performed data analysis, generated figures, wrote manuscript. J.C., R.X., M.H., W.L., performed experiments and X.Z., performed data process. All authors contributed to edit manuscript.

## Data availability

scRNAseq data sets have been deposited in the Gene Expression Omnibus (GEO) database under the accession code: GSE210669.

