## Supplementary material for "Intestinal stem cell aging at single-cell resolution: functional perturbations alter cell developmental trajectory reversed by gerotherapeutics": included

Figure S1

a

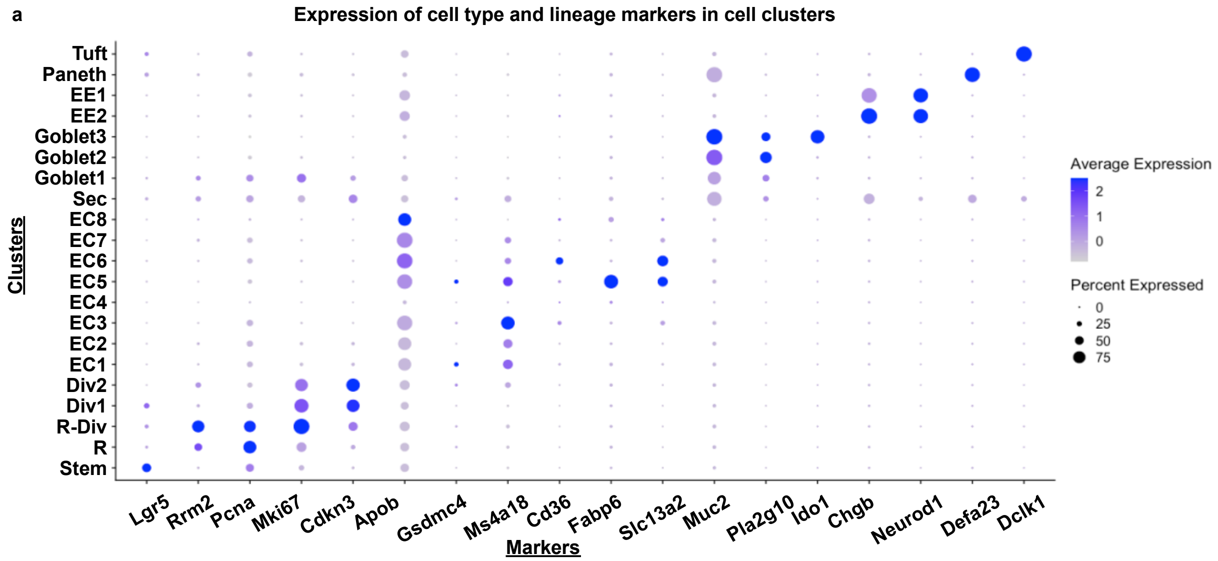


b

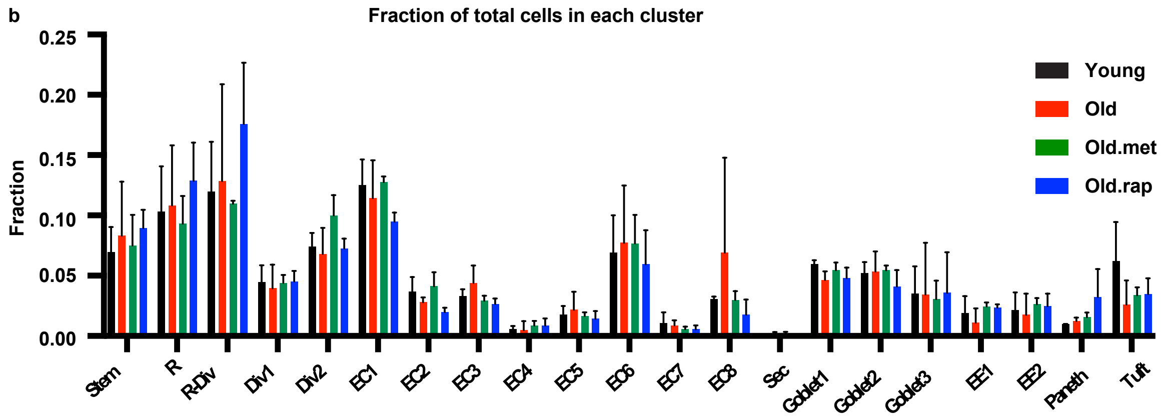


**Figure S1. scRNAseq analysis of small intestinal epithelial cells: a,** dotplot showing marker genes used to define cell type of each cluster identified by Seurat. **b,** fraction of each cluster in each condition: mean value with standard deviation is plotted.

Figure S2

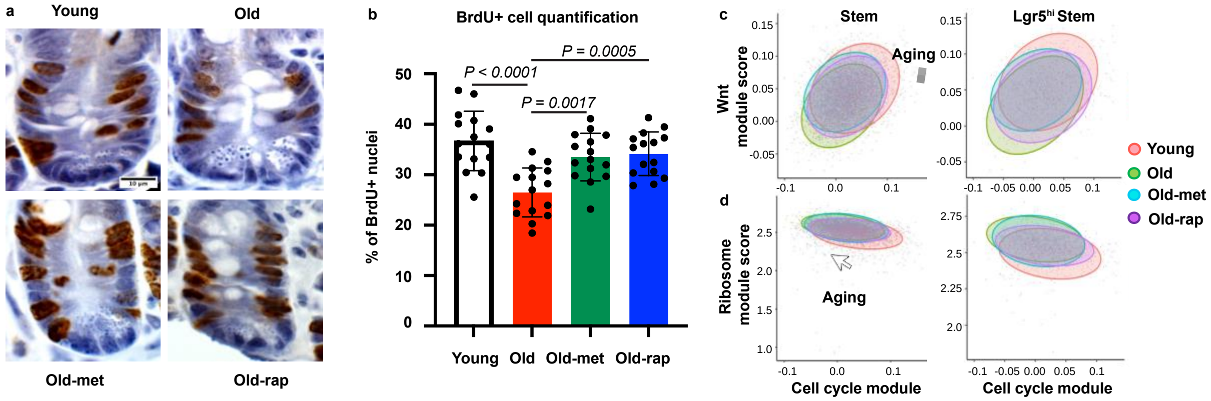


**Figure S2. Wnt and cell cycle is coordinated in individual cells, but Ribosome and cell cycle are not: a,** Representative images of BrdU staining on fixed tissue. Scale bar: 10 μm. **b,** quantification of BrdU+ nuclei and statistical analysis using ANOVA followed by multiple comparison (15 duodenal crypts were analyzed from 3 mice of each condition) **c-d,** Correlated distribution of activities of Wnt pathway and cell cycle pathway in Stem cluster or Lgr5$^{hi}$ ISCs (c) or correlated distribution of activities of Ribosome pathway and cell cycle pathways in Stem cluster or Lgr5$^{hi}$ ISCs (d). Colored ellipse shows the distribution range of 95% of total cells.

Figure S3

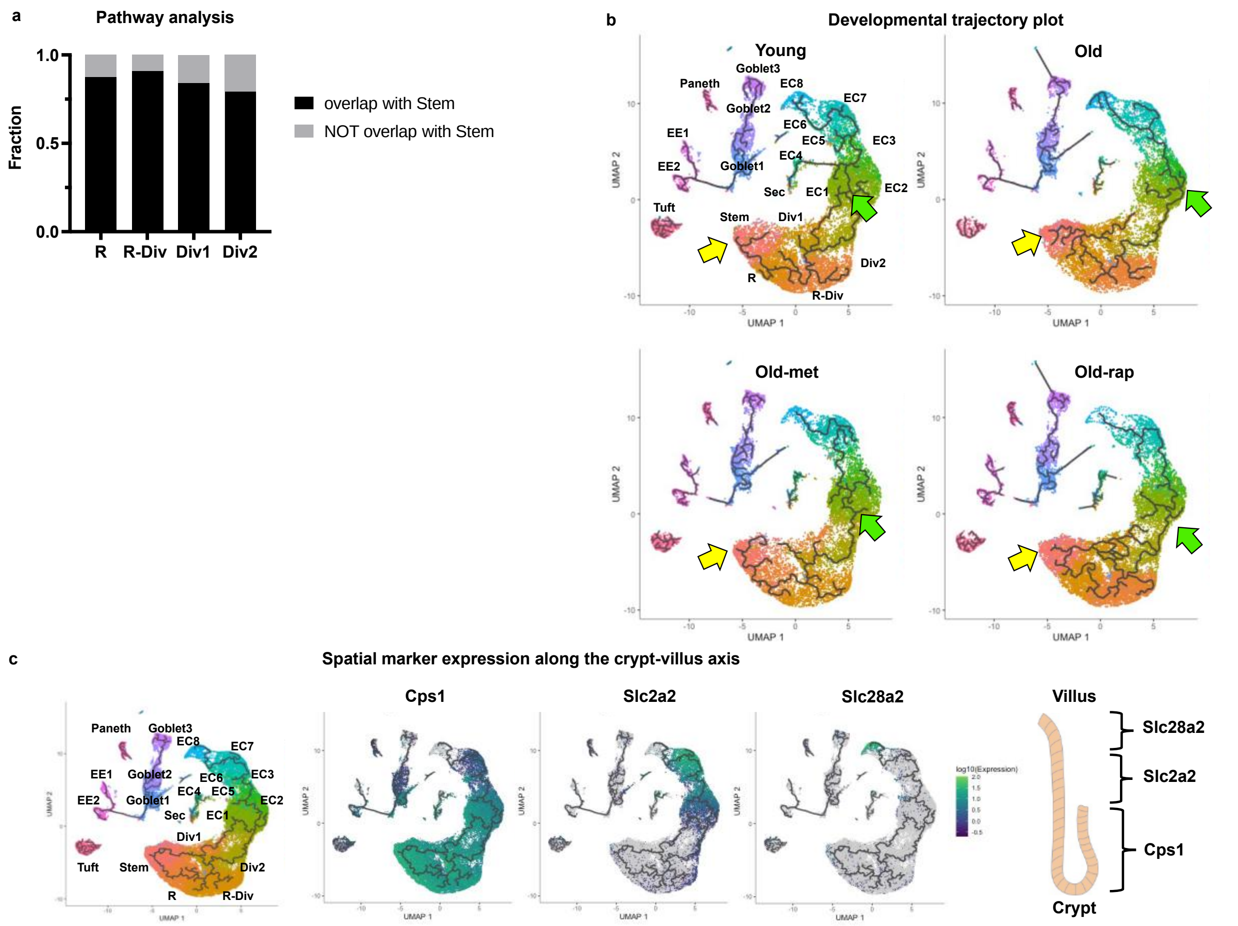


**Figure S3. Impact of aging on stem cells as a function of position along the developmental trajectory: a,** barplot showing overlapping fraction of pathway analysis in progenitor cell compartments with Stem cluster. **b,** trajectory plot of each condition with cluster identified from Seurat. Yellow arrow indicates the Stem cluster as a start point and green arrow indicates transition point from progenitor cell compartment to enterocyte compartment. **c,** trajectory plot with expression level of location-specific genes, adopted from (Moor et al., 2018).

### Table S1. Number of differentially expressed genes

| celltype | differential genes O vs Y | reversed by Met | reversed by Rap | reversed by both |
|---|---|---|---|---|
| Stem | 68 | 21 | 15 | 11 |
| R | 62 | 25 | 10 | 9 |
| R_Div | 80 | 28 | 11 | 9 |
| Div1 | 69 | 24 | 12 | 8 |
| Div2 | 58 | 21 | 7 | 7 |
| EC1 | 82 | 15 | 7 | 6 |
| EC2 | 76 | 16 | 16 | 10 |
| EC3 | 159 | 55 | 19 | 12 |
| EC4 | 8 | 2 | 2 | 2 |
| EC5 | 30 | 10 | 8 | 5 |
| EC6 | 94 | 39 | 32 | 18 |
| EC7 | 78 | 10 | 5 | 0 |
| EC8 | 564 | 431 | 399 | 350 |
| Sec | 2 | 0 | 2 | 0 |
| Goblet1 | 57 | 12 | 3 | 3 |
| Goblet2 | 31 | 3 | 3 | 1 |
| Goblet3 | 40 | 14 | 20 | 11 |
| EE1 | 6 | 1 | 2 | 1 |
| EE2 | 9 | 1 | 2 | 1 |
| Paneth | 31 | 5 | 1 | 0 |
| Tuft | 76 | 2 | 1 | 0 |

**Table S2. Multiple comparison for Figure 3b-i**

| | | Significance | | | |
|---|---|---|---|---|---|
| | | Wnt | Cell cycle | Ribosome | OXPHOS |
| main | Y vs O | **** | **** | * | ** |
| | Y vs O-met | ns | ** | ns | ns |
| | Y vs O-rap | *** | ** | ns | ** |
| | O vs O-met | *** | ** | ns | * |
| | O vs O-rap | ns | ** | ns | ns |
| | O-met vs O-rap | ** | ns | ns | * |
| side | Y vs O | ns | ns | ns | ns |
| | Y vs O-met | ns | * | ns | ns |
| | Y vs O-rap | ns | ns | ns | ns |
| | O vs O-met | ns | ns | ns | ns |
| | O vs O-rap | ns | ns | ns | ns |
| | O-met vs O-rap | ns | ns | ns | ** |

$P < 0.0001$ ****
$P < 0.001$ ***
$P < 0.01$ **
$P < 0.05$ *
